# Exploration of miRNA-mediated fertility regulation network of cytoplasmic male sterility during flower bud development in soybean

**DOI:** 10.1101/351163

**Authors:** Ding Xianlong, Zhang Hao, Ruan Hui, Li Yanwei, Chen Linfeng, Wang Tanliu, Jin Ling, Li Xiaoqiang, Yang Shouping, Gai Junyi

**Affiliations:** Soybean Research Institute, National Center for Soybean Improvement, Key Laboratory of Biology and Genetic Improvement of Soybean (General, Ministry of Agriculture), State Key Laboratory of Crop Genetics and Germplasm Enhancement, Jiangsu Collaborative Innovation Center for Modern Crop Production, College of Agriculture, Nanjing Agricultural University, Nanjing 210095, China; College of Life Sciences, Nanjing Agricultural University, Nanjing 210095, China

**Keywords:** Soybean (*Glycine max* (L.) Merr.), Cytoplasmic male sterility, Flower bud, miRNA, Target gene

## Abstract

Cytoplasmic male sterility (CMS) plays an important role in the production of soybean hybrid seeds. MicroRNAs (miRNAs) are a class of non-coding endogenous ~21 nt small RNAs that play crucial roles in flower and pollen development by targeting genes in plants. Here, two small RNA libraries and two degradome libraries were constructed from the flower buds of the soybean CMS line NJCMS1A and its restorer (Rf) line NJCMS1C. Following high-throughput sequencing, 558 known miRNAs, 103 novel miRNAs on the other arm of known pre-miRNAs, 10 novel miRNAs, and a number of base-edited miRNAs were identified. Among the identified miRNAs, 76 differentially expressed miRNAs were discovered with greater than two-fold changes between NJCMS1A and NJCMS1C. By degradome analysis, a total of 466 distinct transcripts targeted by 200 miRNAs and 122 distinct transcripts targeted by 307 base-edited miRNAs were detected. Further integrated analysis of transcriptome and small RNA found some miRNAs and their targets’ expression patterns showing a negative correlation, such as miR156b-GmSPL and miR4413b-GmPPR. Previous reports showed that these targets might be related to flower bud development, suggesting that miRNAs might act as regulators of soybean CMS fertility. These findings may provide a better understanding of the miRNA-mediated regulatory networks in CMS mechanisms of soybean.

## INTRODUCTION

Soybean is one of the most important oil crops in the world. However, soybean yields are relatively low and the utilization of heterosis is one of the effective ways to improve them. Cytoplasmic male sterility (CMS) is a simple and efficient pollination control system that plays an important role in the production of hybrid seed. CMS is caused by mitochondrial genes with coupled nuclear genes, and CMS-based hybrid seed technology uses a three-line system: the CMS line, the maintainer line, and the restorer of fertility (Rf) line (Chen *et al.* 2014). The CMS line is propagated by crossing with the maintainer line and by crossing the CMS line with the Rf line, the male fertility of the Fi plants can be restored by the Rf gene(s) that come from the nuclear genome of the Rf line. Whether it is the production of CMS or F_1_ fertility restoration, a series of changes will occur at both the molecular and physiological levels. In this process, the interaction between genes is very obvious. Therefore, the molecular mechanism of CMS is a hot research topic and there may be a high degree of epigenetic regulation involvement.

MicroRNAs (miRNAs) are a class of non-coding endogenous ~21 nt small RNAs (Bartel *et al.* 2004) that play vital regulatory roles in plant growth and development, such as flowering-time regulation (Nie *et al.* 2015), heat tolerance (Liu *et al.* 2017a), accumulation of anthocyanins (Liu *et al.* 2017b), response to the phytohormone abscisic acid (Duan *et al.* 2016), cotton fiber elongation (Wang *et al.* 2016b), pollen or flower bud development (Shen *et al.* 2011; Yang *et al.* 2013; Jiang *et al.* 2014; Ding *et al.* 2016; Zhang *et al.* 2016), and so on. Among all of the miRNA families, MIR156 is one of the most conserved families in plants (Wang *et al.* 2016c) and regulates the vegetative phase change floral transition, integrated into flowering pathways by target squamosa promoter binding protein-like (SPL) transcription factors (Spanudakis *et al.* 2014). Moreover, the miR156 targeted or non-targeted SPL genes function together to redundantly secure male fertility in *Arabidopsis* (Xing *et al.* 2010). Compared with conserved miRNA, species-specific miRNA expression is relatively low, and they can be induced to function under certain special conditions (Tang, 2010).

In this study, a number of miRNAs and their targets in soybean were identified by comparing a CMS line (NJCMS1A) and its Rf line (NJCMS1C), based on small RNA sequencing and degradome analysis. The results may improve our understanding of the regulatory role of miRNAs in controlling soybean CMS fertility.

## MATERIALS AND METHODS

### Plant materials, sample collection, and total RNA extraction

The soybean cytoplasmic male-sterile line NJCMS1A was developed through consecutive backcross procedures with the cultivar N8855 as donor parent and N2899 (hereafter designated as NJCMS1B) as recurrent parent (Gai *et al.* 1995; Ding *et al.* 1999; Ding *et al.* 2002). “Zhongdou 5” (hereafter designated as NJCMS1C for further evaluation of the genetic mechanism of male fertility restoration) was identified as the restorer of NJCMS1A in a previous study (Yang *et al.* 2007). NJCMS1A and NJCMS1C were planted in the summer of 2013 at Jiangpu Experimental Station, National Center for Soybean Improvement, Nanjing Agricultural University, Nanjing, Jiangsu, China, for sample collection for small RNA sequencing and degradome sequencing. The male-sterile plants and fertile plants were identified through three methods, namely, the dehiscence of anthers, germination rate of pollen, and performance of plants at maturity. Cytological observation showed that the male abortion of NJCMS1A occurred mainly at the early binucleate pollen stage (Fan, 2003).

Because it is very difficult to judge the precise developmental stage of pollen from the appearance of the flower buds in soybean, during the flowering period in the summer of 2013, flower buds of different sizes up to and including the abortion stage were collected and mixed from NJCMS1A and NJCMS1C plants, and then immediately frozen in liquid nitrogen and stored at −80°C for further use. Flower buds were collected from NJCMS1A and NJCMS1C in the summer of 2015 for Quantitative real-time PCR (qRT-PCR). Total RNAs from the flower buds of NJCMS1A and NJCMS1C were extracted using TRIzol reagent (Invitrogen, Carlsbad, CA, USA) according to the manufacturer’s protocol.

### Small RNA sequencing library construction and bioinformatics analysis

Two small RNA libraries were constructed and sequenced by the Beijing Genomics Institute (BGI, Shenzhen, Guangdong Province, China) using the Illumina HiSeq 2000 System (TruSeq SBS KIT-HS V3, Illumina, Santa Clara, CA, USA). After sequencing, raw sequencing reads were processed into clean reads by filtering out adaptor contaminants, oversized insertions, low-quality reads, poly(A) tags, and small tags (<18 nt). The small RNA tags were mapped to the soybean genome (Wm82.a2.v1) using SOAP to analyze their expression and distribution on the genome. All small RNA sequences were used to query the RNA sequences for deposited repeats (RepeatMasker output) in GenBank (NCBI GenBank). Rfam (V 11.0) was used to separate the small RNAs that matched rRNA, scRNA, snoRNA, snRNA, and tRNA and remove the degraded mRNA fragments (Wm82.a2.v1) in the small RNA tags that aligned to exons and introns. Next, to identify conserved miRNAs, clean reads were compared with known miRNAs of soybean deposited at miRBase 21.0 (http://www.mirbase.org/). Finally, the other sequences that did not map to known miRNAs and other kinds of small RNA were referred to as unannotated sequences for identification of new members of known miRNA families and for novel miRNA prediction.

Potentially novel miRNAs were identified by using MIREAP (https://source-forge.net/projects/mireap/), and their secondary structures were predicted by the mfold web server (http://mfold.rna.albany.edu/?q=mfold/RNA-Folding-Form) (Zuker, 2003). The criteria used for selecting novel miRNAs must meet the following characteristics based on Meyers *et al.* (2008), Zhang *et al.* (2006), Kozomara *et al.* (2014), and miRBase 21.0: (1) The candidate miRNA-5p and miRNA-3p are derived from opposite stem arms with minimal matched nucleotide pairs exceeding 16 nt and with maximal size differences of up to 4 nt; (2) The most abundant reads from each arm of the precursor must pair in the mature miRNA duplex with a 2-nt 3′overhang; (3) The number of asymmetric bulges within the miRNA-5p/miRNA-3p duplex must be one or fewer, and the size of the asymmetric bulges must be two bases or smaller; (4) The miRNA-5p and miRNA-3p are simultaneously on the two arms of pre-miRNA secondary structures, and one of the mature miRNAs must be no fewer than five; (5) The candidate miRNA precursor must have high negative minimum free energy (MFE) and minimum folding energy index (MFEI), with MFE <−0.2 kcal/mol/nt and MFEI > 0.85.

### Degradome library construction, data analysis, and target identification

Two degradome libraries were constructed according to the methods of Addo-Quaye *et al.* (2008) and German *et al.* (2008) and sequenced on the Illumina HiSeq 2000 (TruSeq SBS KIT-HS V3, Illumina, BGI, Shenzhen, Guangdong Province, China). Alignments with scores up to 4.5, where G:U pairs scored 0.5 and no mismatches were found at the site between the 10th and 11th nucleotides of the corresponding miRNAs, were considered potential targets.

### Quantitative real-time PCR validation

Stem-loop qRT-PCR and qRT-PCR were carried out to validate differential expression levels of miRNAs and miRNA targets, respectively (Chen *et al.* 2005). All primers (Table S1) were designed based on the mature miRNA and mRNA sequences and synthesized commercially (Invitrogen, Shanghai, China). According to the procedures provided in the iScript Select cDNA Synthesis Kit (containing GSP enhancer solution, BIO-RAD, USA), 1 μg of total RNA was reverse-transcribed by iScript reverse transcriptase using stem-loop primers. The qRT-PCR analysis was carried out using iTaq Universal SYBR Green Supermix (BIO-RAD, USA) on a Bio-Rad CFX96 instrument (CFX96 Touch, BIO-RAD, USA). All reactions were run with three biological replicates, and miR1520d (Kulcheski *et al.* 2010) was used as the internal control gene. The relative expression levels of miRNAs and genes were quantified using the 2^−ΔΔCt^ method. Student’s *t* test was performed to compare miRNA or mRNA differences in expression between NJCMS1A and NJCMS1C. Expression levels that significantly differed (*p*<0.05) according to Student’s *t* test were labeled as: “*”, “**”, and “***”, which represent *p*<0.05, *p*<0.01, and *p*<0.001, respectively, which indicated significant difference levels between NJCMS1A and NJCMS1C.

### Data availability

All of the small RNA-seq and degradome-seq data were submitted to the National Center for Biotechnology Information (NCBI) under the accession number PRJNA304685 and --.

## RESULTS

### Overview of small RNA library data sets

Two small RNA libraries were constructed using total RNA obtained from flower buds of the CMS line NJCMS1A and its Rf line NJCMS1C and sequenced by Illumina Solexa high-throughput sequencing technology. After removing adaptor contaminants and low-quality reads etc., a total of 11,974,839 and 12,072,909 clean reads were obtained from the two libraries, respectively, ranging from 18-nt to 30-nt in length (Figure 1A). Using the SOAP software, more than 80% small RNA tags were perfectly mapped to the soybean genome (Table 1).

**Figure 1.**
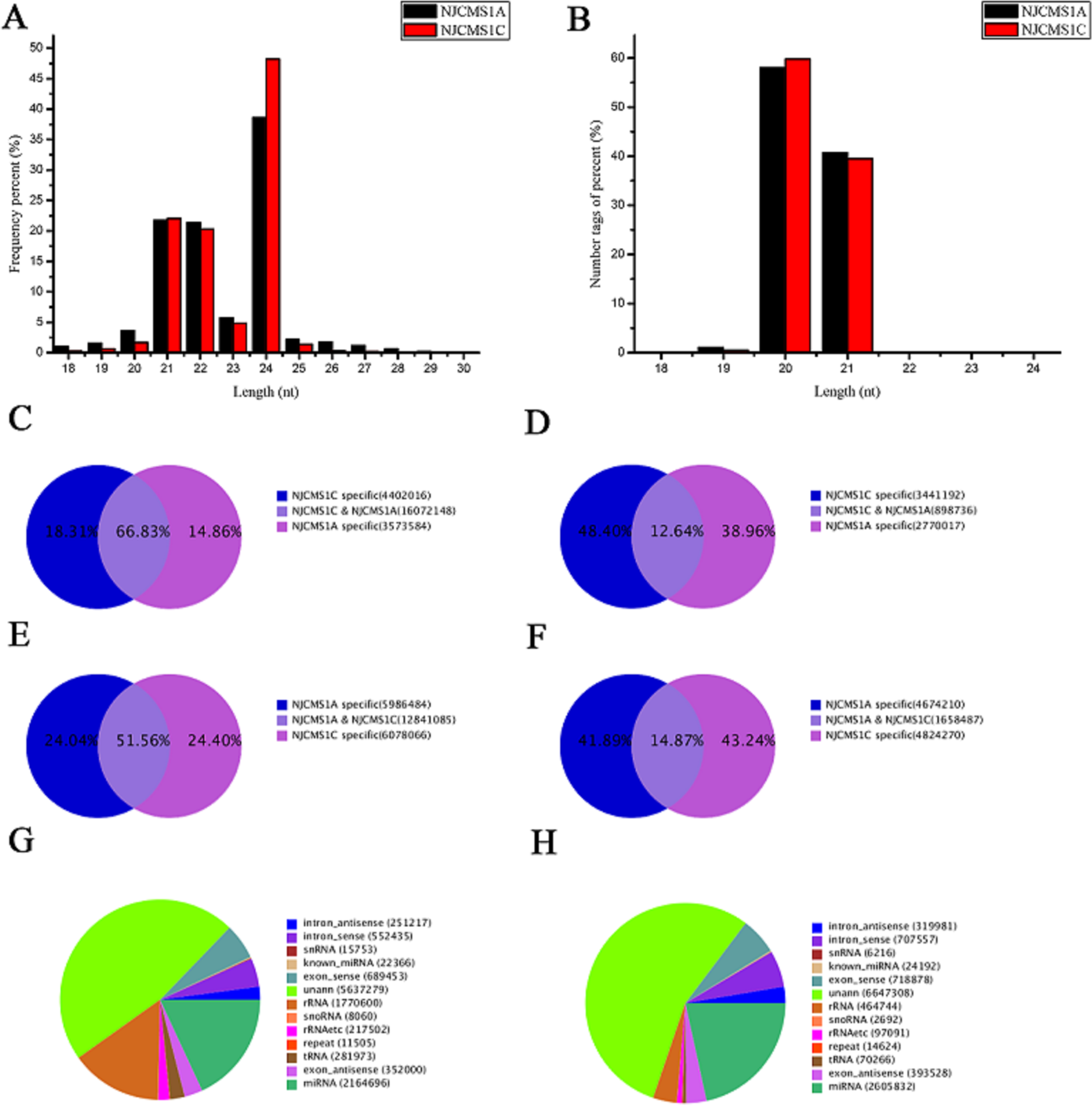
Distribution of small RNAs and degradome-seq clean tags. **A-B:** Length distribution of small RNAs and degradome-seq tags. **C-D:** Total sRNAs and unique sRNAs in small RNA libraries. **E-F:** Total tags and unique tags in degradome libraries. **G-H:** Small RNA annotation in NJCMS1A and NJCMS1C.

**Table 1.**
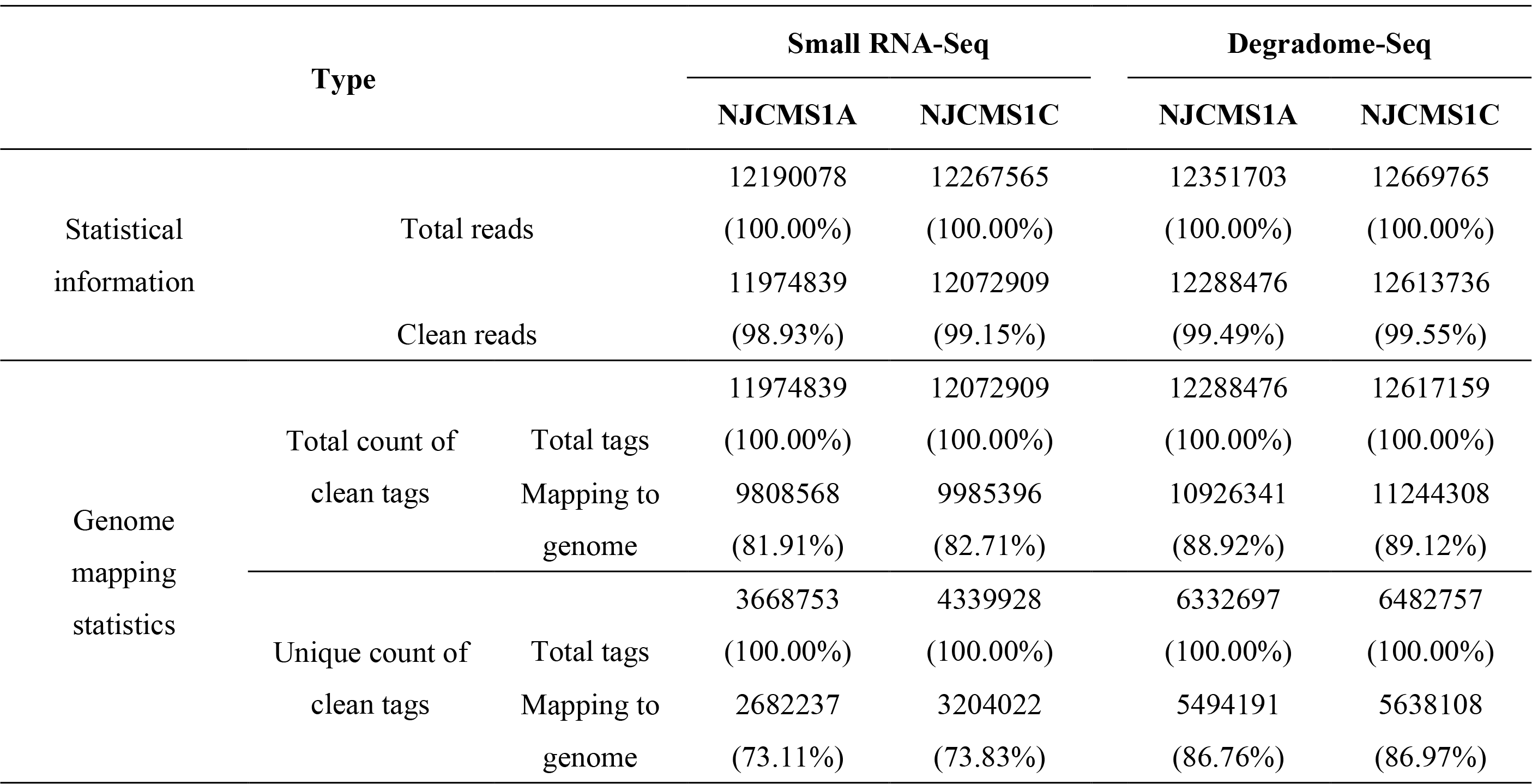
Summary of small RNA sequences and degradome sequences.

Upon analyzing the small RNA tags between NJCMS1A and NJCMS1C, we found that NJCMS1A and NJCMS1C comprised 14.86% and 18.31% of the total sRNAs (Figure 1C), respectively. Only 12.64% of unique sRNAs were shared between the CMS line and its Rf line (Figure 1D). Moreover, the NJCMS1C-specific unique sRNAs were greater than those of NJCMS1A, which means that most of the unique sRNAs found in NJCMS1C flower buds were different from those in NJCMS1A (Figure 1C, D).The small RNAs annotation for the two samples is shown in Figure 1g and h. The majority of small RNAs were 21–24 nt, and 24-nt small RNAs were the most abundant (Figure 1A), which is in accordance with the previously reported miRNAs of soybean (Song *et al.* 2011).

### Identification of known miRNAs and novel miRNAs on the other arm of known pre-miRNAs in soybean

The mappable small RNA sequences were aligned with the known soybean miRNAs using miRBase 21.0. A total of 499 known pre-miRNAs corresponding to 558 mature miRNAs that belong to 222 families were detected (Table S2, S3). The sequences of some miRNAs were tagged with “D” (Table S2), representing that these were either shorter or longer than the mature miRNAs in miRBase 21.0; these can be classified as isomiRNAs and may share the same target genes with the mature miRNAs. Through high-throughput sequencing, 103 novel miRNAs on the other arm of known pre-miRNAs were identified in two samples (Table S4). The predicted secondary structure of MIR160f was selected as an example and is shown in Figure S1A.

### Identification of novel miRNAs

To identify novel miRNAs, all unannotated and mappable small RNAs were aligned to known plant miRNAs in miRBase 21.0. The small RNAs that could not be mapped to known plant miRNAs were classified as novel miRNAs. The five criteria described in the materials and methods section were utilized to increase predictive accuracy. As a result, ten novel miRNAs (five pairs) that belong to four families were identified in this study (Table 2, Table S3, S5). Precursors of these novel miRNAs were identified by MIREAP and varied from 85 to 93 nt in length, with MFE values ranging from −0.47 to −0.57 kcal/mol/nt. The MFEI values ranged from 1.15 to 1.65 (Table S5), all of them being greater than 0.97 and the average being 1.36. The length of all of the mature miRNAs was 21 nt. Some of the novel miRNAs, such as novel_mir_003a to novel_mir_003b, shared the same mature sequence but had different precursors that came from different loci. The secondary structure for the precursor of novel_mir_003a was selected as an example and is shown in Figure S1B.

**Table 2.**
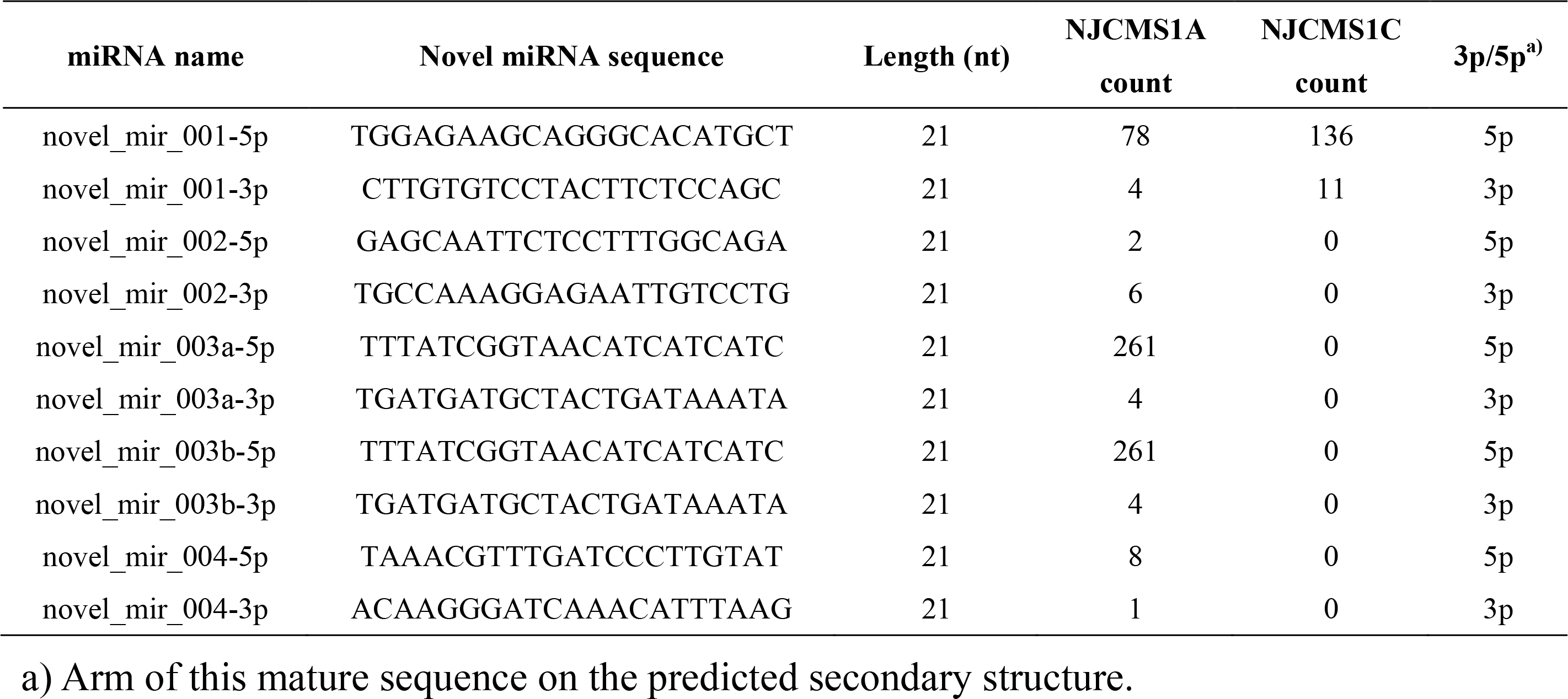
Novel miRNAs identified in NJCMS1A and NJCMS1C.

### Overview of miRNA with base edits

In the present study, we detected miRNA with base edits by aligning unannotated sRNA tags with mature miRNAs from miRBase21.0. A total of 174 miRNA members exhibited base edits and the miRNA editing frequency was 0.13% to100.00% (Table S6-1, Table S6-2). Some miRNAs, such as miR1516a-3p, were not found by sequencing, but its base-edited miRNA was detected in this study (Table S6-1).The miRNA base edits had truly occurred in soybean, as confirmed by our sequencing results (Figure S2). Most of the miRNA editing events occurred at nucleotide positions of 5-17 and the miRNA editing patterns were similar between the two lines, but the editing frequency at position of 11 is slightly lower (Figure 2A). The most dominant nucleotide substitution types were A->G, C->U and G->U (Figure 2B).

To validate the occurrence of the miRNA editing, the two most frequently observed editing types (A->G and C->U) were examined. The precursor miRNA sequence was cloned from the soybean genome, and the mature miRNA sequence was cloned from the cDNA reverse transcribed by the stem-loop reverse primers. Both sequencing and validate results confirmed that nucleotide substitutions had truly occurred during the mature miRNA processing (Table S6, Figure 2C, D). However, the reason for the variant edition type and their function in mature miRNA processing or gene expression regulation remains elusive and warrants further study.

**Figure 2.**
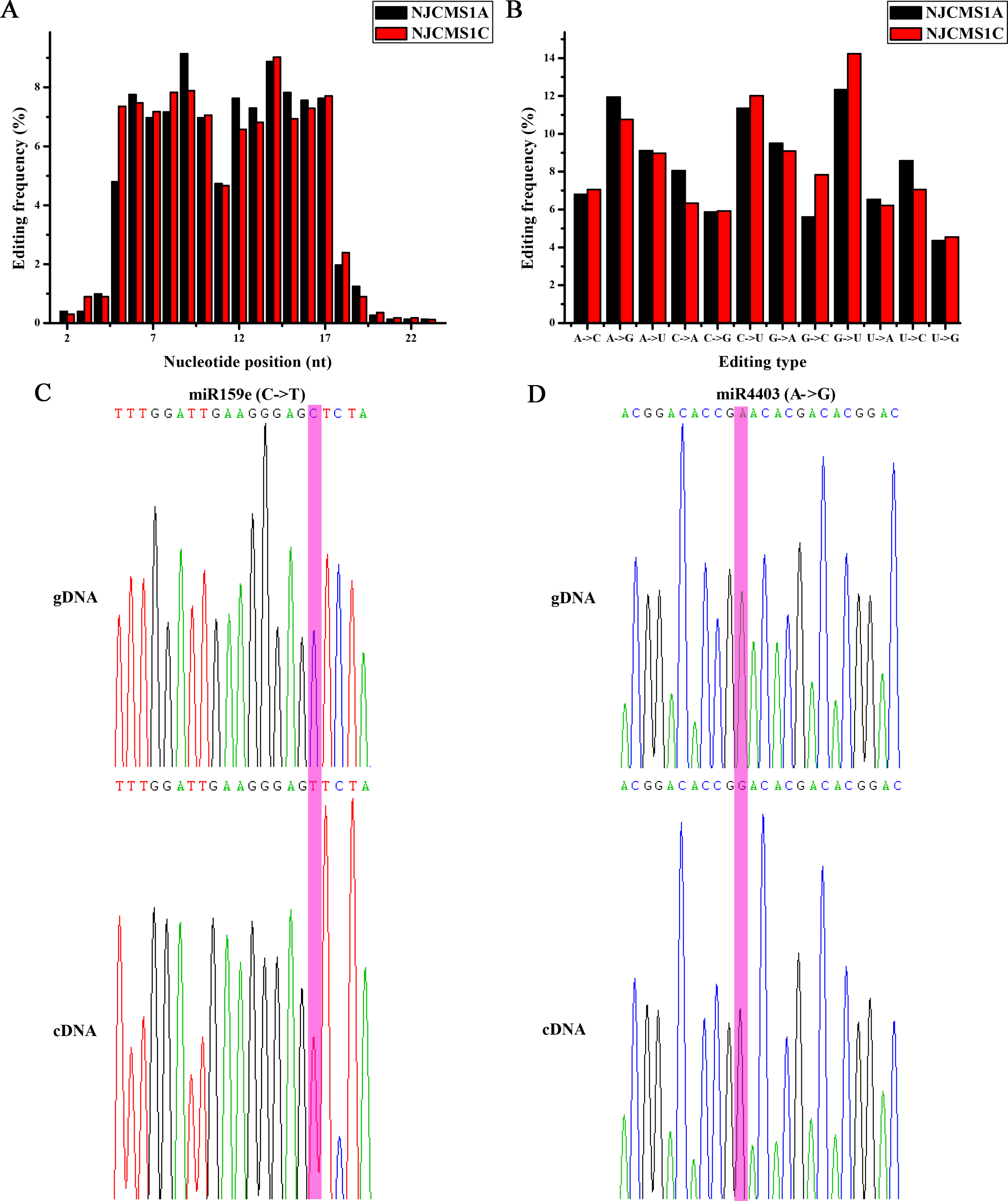
Analysis of miRNA editing events. **A:** miRNA editing frequency at each miRNA nucleotide position in both NJCMS1A and NJCMS1C. **B:** miRNA editing types and their frequency in both NJCMS1A and NJCMS1C. **C-D:** Validation of miRNA editing types inferred from highthroughput sequencing by precursor miRNA and mature miRNA cloning and sequencing. Sequencing chromatograms from the two observed editing types (C->T, A->G) of miR159e and miR4403 are shown. The edited positions are highlighted inpink. The upper panel indicates the part of precursor miRNA cloned from genomic DNA, and the lower panel indicates the mature miRNA cloned from cDNA reverse transcribed with stem-loop RT primers. Primers are listed in Table S1.

### Comparative analysis of small RNAs

Throughout all the identified miRNAs in the flower buds of NJCMS1A and NJCMS1C, 76 differentially expressed miRNAs (including 74 known miRNAs and two novel miRNAs) with more than two-fold relative change was identified (Table S7). Among the differentially expressed miRNAs, 28 miRNAs were up-regulated in NJCMS1A, and the remaining miRNAs showed higher expression levels or were only expressed in the flower buds of NJCMS1C (Table S7). The miR4393a and miR5667 were only expressed in NJCMS1A, and miR4346, miR4372a, miR4400, miR4994-5p, and miR5785 were only detected in NJCMS1C. In particular, gma-miR4346 showed a high relative expression level and may have a unique function in the process of pollen development in NJCMS1C. Of the differentially expressed novel miRNAs, most of them were only expressed in NJCMS1A or NJCMS1C (Table S7).

To examine miRNA expression, ten selected differentially expressed miRNAs were assayed by stem–loop qRT-PCR. As showed in Figure 3, nine of these were consistent with the sequencing reads. However, miR6300 was found to be up-regulated in NJCMS1C, but was enriched in NJCMS1A based on the high-throughput sequencing analysis and this result warrants further study.

**Figure 3.**
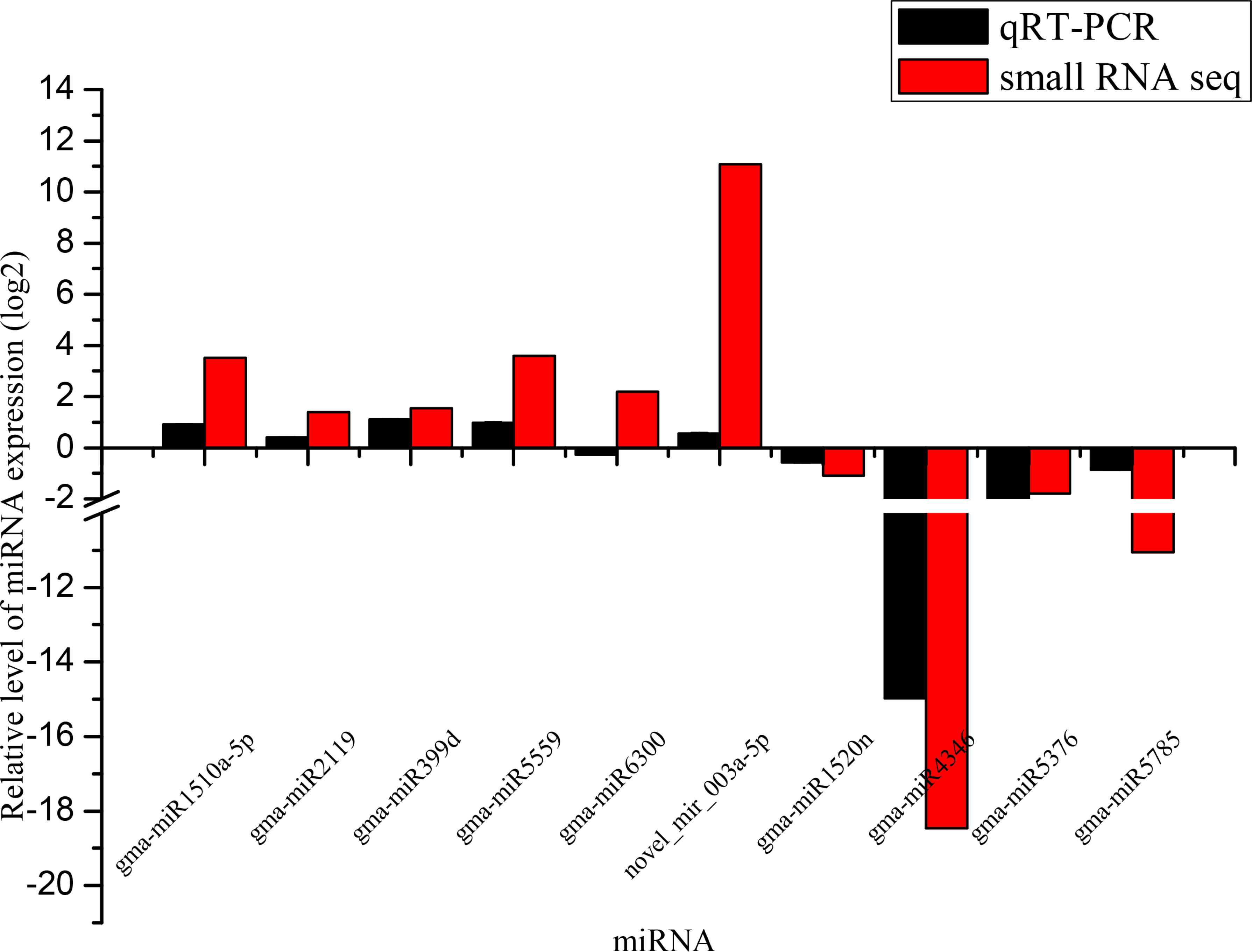
Verification results of differentialy expressed miRNAs between NJCMS1A and NJCMS1C. The *y* axis indicates the miRNA relative expression level (log2) generated from qRT-PCR analysis and high-throughput sequencing. The results were obtained from three biological replicates, and the error bars indicate the standard error of the mean of 2^−ΔΔCt^, with NJCMS1C as a control.

**Figure 4.**
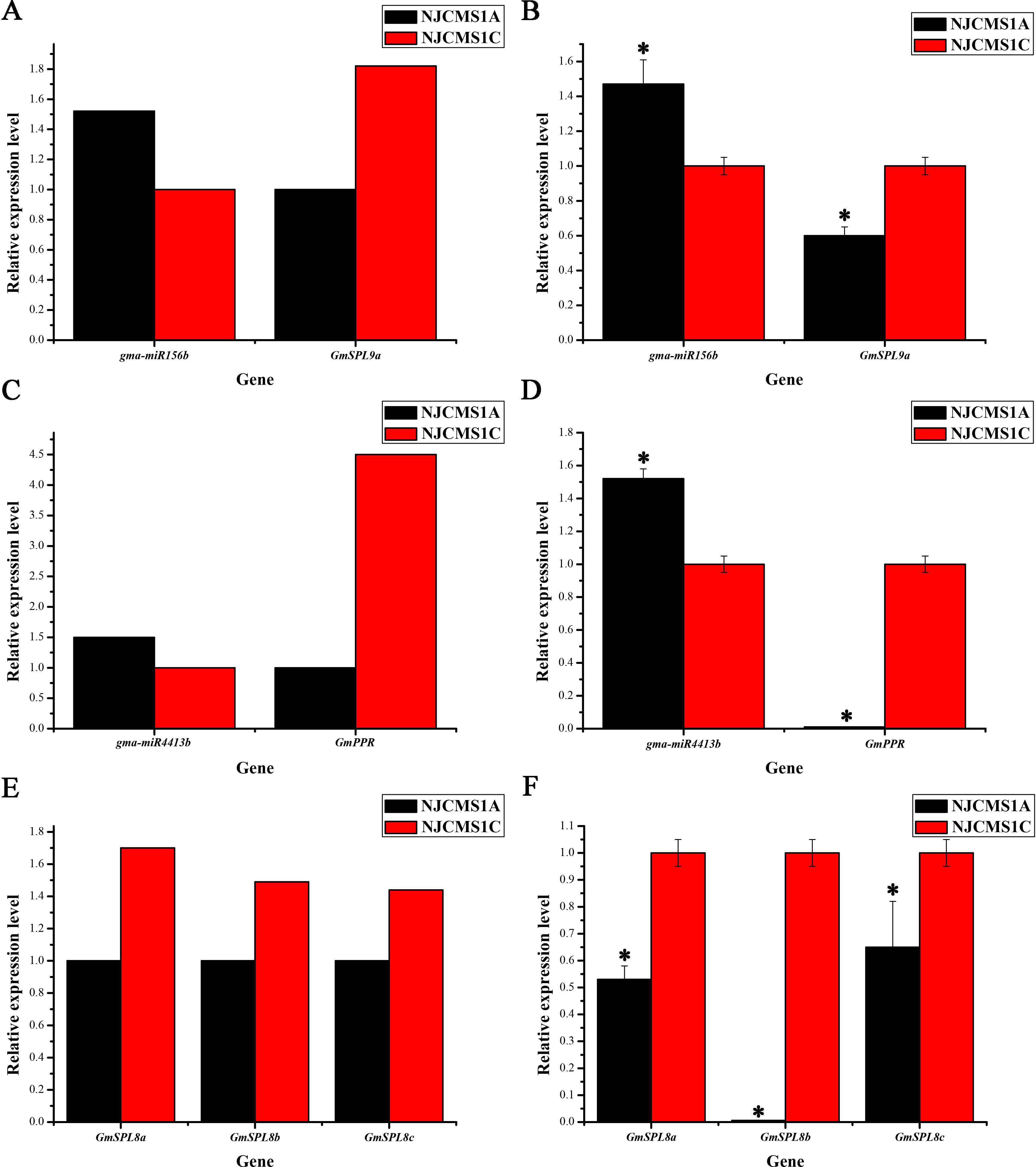
Relative expression level of miRNA and its targets between NJCMS1A and NJCMS1C. **A, C and E:** Relative expression level of miRNA and its targets in small RNA sequencing and transcriptome analysis, respectively. **B, D and F:** Relative expression level of miRNA and its targets in qRT-PCR analysis.

### Degradome library construction and comparative analysis of miRNA targets between NJCMS1A and NJCMS1C

Here, we performed degradome sequencing and analysis to search for the miRNA targets in NJCMS1A and NJCMS1C. A total of 12,288,476 and 12,617,159 clean reads were generated from the NJCMS1A and NJCMS1C libraries, respectively (Table 1). More than 98.00% of the tags were 20 or 21 nt in length (Figure 1B), as expected from the normal length distribution peak of degradome fragments. Of the unique tags, 41.89% and 43.24% were from NJCMS1A and NJCMS1C, respectively, and only 14.87% of them were shared between NJCMS1A and NJCMS1C (Figure1E, F).

To better understand the function of miRNA in flower bud development, we performed a conjoint analysis of the miRNA and degradome together. A total of 466 distinct transcripts targeted by 200 miRNAs were detected (Table S8, Figure S3). These transcripts included SPL family protein, GAMYB protein, auxin response factor, homeobox-leucine zipper protein, nuclear transcription factor Y, GRAS family transcription factor, and transcription factor APETALA2, which have essential roles in gene regulation (Table S8). In addition, a set of pentatricopeptide-repeat-containing (PPR) proteins were targeted by miR1508, miR160, miR171o, miR4413b and miR5674, which may be a group of Rf proteins that suppress CMS gene expression through posttranscriptional mechanisms such as editing, cleavage, and degradation of the target mRNAs (Table S9, Chen *et al.* 2014).

To determine if the base edit in positions 2–8 and 10–11 of a mature miRNA can change its target recognition, a combined analysis of the degradome library and base-edited miRNAs was conducted. As a result, 122 distinct transcripts targeted by 307 base-edited miRNAs were detected (Table S10). Most of the targets of the base-edited miRNAs were the same as for the corresponding miRNAs for which no base editing had occurred (Table S11). However, some targets were filtered out by the base editing. For example, there were 75 targets of miR156b and miR156f, but as a result of the base edited, only 61 targets were detected (Table S11). Interestingly, we also detected some new targets that were not targeted by miRNA when no base editing occurred, such as Glyma.05G230700.2 to Glyma.05G230700.4, which were targeted by miR166h-P8CT, miR166k-P8CT, and miR166u-P8CT (Table S11), but not by miR166h, miR166k, and miR166u. Furthermore, with base editing in position 10-11 of the mature miRNA, some miRNA targets were changed. For example, no targets were detected for gma-miR4403 in this study, but as a result of a base edit at position 11, one target (Glyma.17G050600.1) was found (Figure S4, Table S11).

### Integrated analysis of miRNA and mRNA expression between NJCMS1A and NJCMS1C

To better understand the function of differentially expressed miRNAs in flower bud development of soybean, we performed an integrated analysis to identify inverse miRNA-mRNA expression based on small RNA and transcriptome comparative analysis (Li *et al.* (2017)). In consideration of the expression of some miRNAs and their targets were changed more significantly in sporogenic tissues but were submerged by the expression in other tissues of the flower buds, so a cut-off of > 1.5-fold change was used in this section as that of small RNA and transcriptome sequencing in Chinese cabbage CMS (Wei *et al.* 2015). As a result, 13 miRNAs (6 miRNA families) with their 7 inverse expressed target genes that had a fold change >1.5 or <-1.5 were identified (Table S12, Figure S5). Among them, the conserved miRNA, miR156b, was more than 1.5-fold greater in NJCMS1A. Twenty-three target genes were identified to be its targets in this study (Table S8). However, only one target showing a negative correlation with miR156b (Table S12). Despite the conserved miRNAs, miR4392 and miR4413b were two soybean specific miRNAs, which were also altered in NJCMS1A (Table S12). This result indicated species-specific miRNA may also participate in the flower bud development in plant. In addition, 3 conserved miRNA families, miR169, miR1514 and miR397, were more than 1.6-fold greater in the NJCMS1C. Of the targeted genes, 4 were found to be showing more than 2.0-fold higher in NJCMS1A (Table S12).

## DISCUSSION

### Exploration of miRNAs and their targets participating in flower bud development

In plants, many miRNAs seem to be universally expressed across diverse tissues, including the anther. Previous reports have found 100 known miRNAs in the Rf3 and rf3 pollens of maize by Solexa sequencing (Yu *et al.* 2013). In this study, 499 known pre-miRNAs corresponding to 558 mature miRNAs, 103 novel miRNAs on the other arm of known pre-miRNAs, and 10 novel miRNAs were identified in flower buds of the CMS line and its restorer line of soybean. Among all these miRNAs, 76 differentially expressed miRNAs with more than two-fold change between the flower buds of NJCMS1A and NJCMS1C were identified (Table S7). Moreover, miR4393a and miR5667 were only expressed in NJCMS1A and miR4346, miR4372a, miR4400, miR4994-5p, and miR5785 were only detected in NJCMS1C (Table S7); we speculate that these miRNAs might be involved in a specific process of flower bud development. By degradome analysis, 466 targets were chosen and predicted to be cleaved by 200 miRNAs (Table S8). Among the identified targets of miRNAs, some have previously been shown to be involved in floral organ or pollen development, such as miR156 with SPLs (Xing *et al.* 2010), miR172 with APETALA (Aukerman *et al.* 2003), miR159 with GAMYB or GAMYB-like genes (Alonso-Peral *et al.* 2010), and so on.

Moreover, we detected 174 miRNA members in which base editing had occurred (Table S6). Wei *et al.* 2009 divided the miRNA sequence into three major areas; positions 2–8 and 9–16 of a mature miRNA were called the seed region and region A, respectively, and the remainder was denoted region B. miRNA edit events were first found in animals (Wei *et al.* 2009) and subsequently revealed to also occur during plant development (Yan *et al.* 2015). Wei *et al.* 2009 showed that the seed region is used for target recognition; if the seed region and region A or the seed region and region B work together, the miRNA can regulate different species-specific targets. As miRNAs function in plants by either target cleavage or translational repression, editing of the 10th/ 11th positions of miRNA will also lead to altered targets. In order to verify the above results, we combined the analysis of the degradome library and base-edited (seed region and the 10th/11th positions of miRNA) miRNAs. The results showed that most of the targets of base-edited miRNAs were the same as those of the unedited miRNAs (Table S11). However, as the base editing occurred, some targets were filtered. For example, Glyma.08G122000.2 was targeted by gma-miR2118a-3p and gma-miR2118b-3p, but when the base G at position 6 was changed to T, it no longer targeted Glyma.08G122000.2 (Table S11). On the other hand, when the base C at position 11 of gma-miR2118a-3p and gma-miR2118b-3p was changed to T, we identified a new target (Glyma.08G005700.1) by degradome analysis (Table S11).

### Several miRNA regulatory networks might be involved in regulation of flower bud development

It is widely accepted that CMS is caused by mitochondrial genes with coupled nuclear genes, and the male fertility of the F_1_ plants can be restored by the Rf gene(s) that come from the nuclear genome of the Rf line (Chen *et al.* 2014). In this process, the interaction between genes is very obvious, the nuclear genes might change their expression as target genes in response to the mitochondrial signaling, which subsequently affects flower bud development. In this study, altered expression of miRNAs and their targets genes were exhibited between NJCMS1A and NJCMS1C, demonstrating important roles of miRNAs regulation network in the flower bud development of soybean CMS. For example, GmNF-YA3/GmNF-YA5 and GmLAC4 were target genes of miR169 and miR397 (Table S12), respectively. Transcriptome Sequencing showed these target genes were all more than 2-fold change expressed in NJCMS1A (Table S12), and previous studies reported their over-expression affects male gametogenesis and induces nondehiscent anthers in plant (Mu *et al.* 2013; Zhang *et al.* 2014; Nasrin *et al.* 2010). Based on the expression analysis of miRNAs and their corresponding targets, miR156-GmSPL and miR4413-GmPPR regulatory network were selected as two candidate miRNAs-targets combinations that might be involved in flower bud development of soybean CMS.

MIR156 is one of the highly conserved miRNA families in plant, and different members of MIR156 showed highly dynamic expression patterns at different stages of flower development in *Arabidopsis* (Xing *et al.* 2010). Former studies have shown that miR156 plays essential regulatory roles in floral development and male fertility by targeting SPL genes, and the loss of function of miR156/7-targeted SPLs led to a semi-sterile phenotype in *Arabidopsis* (Xing *et al.* 2010). Yu *et al.* 2013 reported that miR156 had a higher expression level in the pollen of S-type CMS in maize. Moreover, it had higher expression levels in the anthers from the three anther developmental stages of the GMS mutant in cotton (Wei *et al.* 2013). In this study, 25 members of MIR156 were found, however, only miR156b and miR156f were more than 1.5-fold change expressed in NJCMS1A (Table S2, S12, Figure S6). And miR156b negatively regulated GmSPL9a that displayed differential expression between NJCMS1A and NJCMS1C, indicating that miR156-SPL regulatory network may participate in flower bud development of soybean (Figure 6). However, a total of 22 target genes were predicted to be targeted by miR156b (Table S8). Among them, 16 target genes were involved in the regulation of transition from vegetative to reproductive phase and flower development through GO annotation (Table S13). However, other genes except GmSPL9a do not reach more than 1.5-fold expression difference between NJCMS1A and NJCMS1C but were highly expressed in flower (Table S12). Previous studies indicated that SPL gene plays a key role in early anther development by promoting other early anther genes required for cell division, specification, and differentiation, resulting in pollen mother cell and tapetum formation (Schiefthaler *et al.* 1999; Yang *et al.* 1999; Xing *et al.* 2010). So we also speculated that some GmSPL genes were also involved in the flower bud development of soybean CMS at the early stage of anther development. In order to verify this view, we used qRT-PCR to measure the expression of miR156b, GmSPL9b and GmSPL13b (random selection) at three periods during flower bud development of soybean. As a result, miR156b was up-regulated at the period I and II, and GmSPL9b and GmSPL13b were down-regulated before period II and III (Figure 5), respectively, which means that the targeted SPL genes exhibit an opposite expression pattern with that of miR156b during certain stage of anther development (Figure 6). Thus, miR156 and SPL may also absolutely required for proper early anther development in soybean CMS.

**Figure 5.**
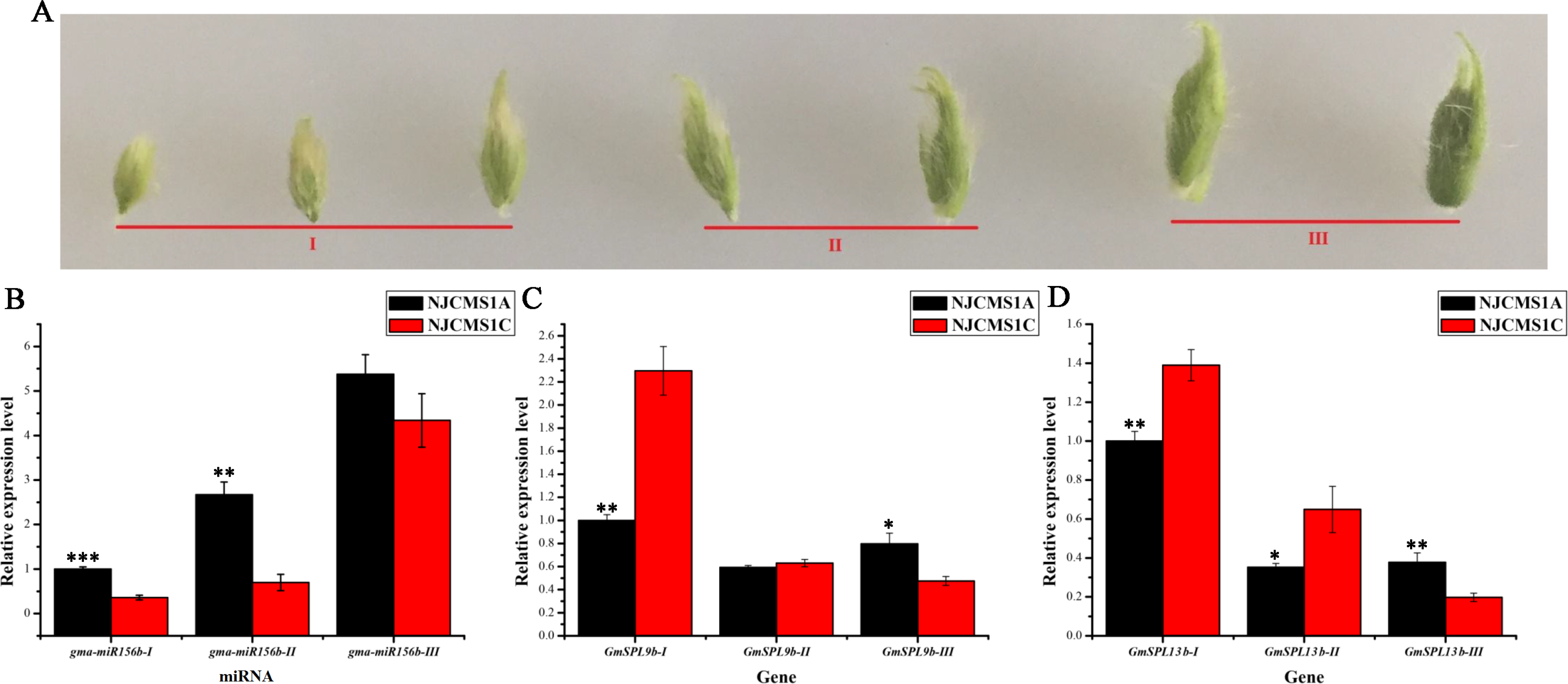
Relative expression level of miR156b and its targets in flower bud of soybean. **A:** the flower bud of three periods that were sampled for qRT-PCR analysis. **B, C and D**: Relative expression level of mi156b, GmSPL9b and GmSPL13b in flower bud of three periods.

**Figure 6.**
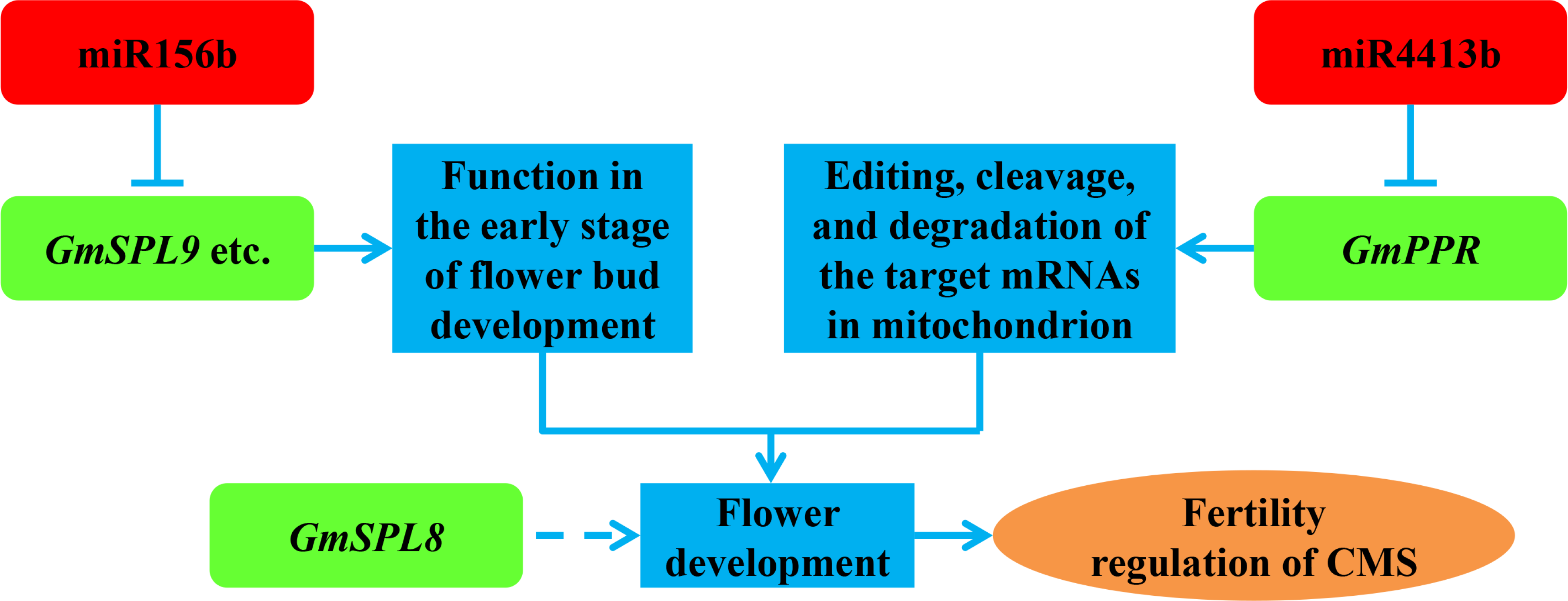
The speculated model of miRNA and its targets regulatory network of soybean CMS during flower bud development. The up-regulated miRNAs and down-regulated mRNAs are in red and green, respectively.

Previous results have also shown that the over-expression of MIR156 in a *spl8* mutant background leads to full male sterility in *Arabidopsis* (Xing *et al.* (2010)) and the introduction of single miR156-resistant SPL transgenes could only partially restore *spl8* mutant fertility. The phenotypic analysis in *Arabidopsis* also showed that a major effect of SPL8 was on microsporogenesis and megasporogenesis within the anthers and ovules, and the absence of a functional SPL8 gene may lead to a failure to enter meiosis (Unte *et al.* (2003)), which means that SPL8 is important to the restoration of pollen fertility. Moreover, SPL8 is not the target gene of miR156 either in *Arabidopsis* (Xing *et al.* (2010)) or in soybean (Table S8). Taking these into account, we achieved the SPL8 expression data in flower bud of NJCMS1A and NJCMS1C by transcriptome comparative and qRT-PCR analysis. The results showed that all of the three GmSPL8 genes were down-regulated in NJCMS1A (Figure 4E, F), and the lower expression of GmSPL8 and up-regulated of miR156b might be one of the reasons for pollen abortion in soybean CMS (Figure 6). All these results indicated that miR156b may have some relationship with GmSPL, as with GmSPL8 and flower development in soybean CMS, which warrants further study.

PPR proteins are classified as Rf proteins that suppress CMS gene expression through posttranscriptional mechanisms such as editing, cleavage, and degradation of the target mRNAs in many plants such as rice, *Brassica,* radish, and sorghum (Chen *et al.* 2014; Brown *et al.* 2003; Hu *et al.* 2012; Iwabuchi *et al.* 1999; Jordan *et al.* 2010; Kazama *et al.* 2003). Recently, Wang *et al.* (2016a) identified the Rf gene for soybean M-type CMS, *Rf-m,* which is located in a PPR gene-rich region on chromosome 16. This region contains 19 putative open reading frames (ORFs), and seven of them encode putative PPR proteins. Two of these (Glyma.16G161900 and Glyma.16G163100) were targeted by miR4413b. miR4413b is an soybean-specific miRNA and was up-regulated (1.5-fold) in NJCMS1A, and its target genes were all PPR proteins (Table S9). Moreover, we found Glyma.16G161900 was significantly down-regulated in NJCMS1A according to the transcriptome data in our lab (Li *et al.* (2017)). Quantitative real-time PCR also showed that miR4413b was up-regulated in NJCMS1A, and Glyma.16G161900 was only expressed in NJCMS1C (Figure 4D). All these results indicated that miR4413b may play a specific function during the formation of soybean CMS (Figure 6), which warrants further study.

## Conclusion

In this study, a large number of miRNAs were identified during flower bud development in the soybean CMS line NJCMS1A and its restorer line NJCMS1C. By deep sequencing, 558 known miRNAs, 103 novel miRNAs on the other arm of known pre-miRNAs, 10 novel miRNAs, and 174 base-edited miRNA members were identified. Among the identified miRNAs, 76 differentially expressed miRNAs were discovered with greater than two-fold changes between NJCMS1A and NJCMS1C. By degradome analysis, a total of 466 distinct transcripts targeted by 200 miRNAs and 122 distinct transcripts targeted by 307 base-edited miRNAs were detected. Small RNA sequencing, transcriptome and qRT-PCR comparative analysis found that miR156-GmSPL-GmSPL8, miR4413-GmPPR regulatory network might be involved in flower bud development of soybean CMS. Further functional studies on these differentially expressed miRNAs will need to provide a better understanding of the miRNA-mediated regulation mechanisms during the CMS occurrence in the soybean N8855-CMS line.

## ACKNOWLEDGMENTS

This work was supported by the National Key Research and Development Program of China (2016YFD0101500, 2016YFD0101504), the National Hightech R&D Program of China (2011AA10A105), and the Program for Changjiang Scholars and Innovative Research Team in University (PCSIRT13073).

## ABBREVIATIONS

AP2: APETALA2
CMS: Cytoplasmic male sterility
GA: gibberellins
MFE: Minimum free energies
MFEI: Minimal folding energy indices
miRNA: microRNA
PPR: Pentatricopeptide repeat-containing
qRT-PCR: Quantitative real time PCR
Rf: restorer
SPL: Squamosa promoter-binding protein-like

## CONFLICT OF INTEREST STATEMENT

The authors declare that they have no competing interests.

---

**Figure S1.**
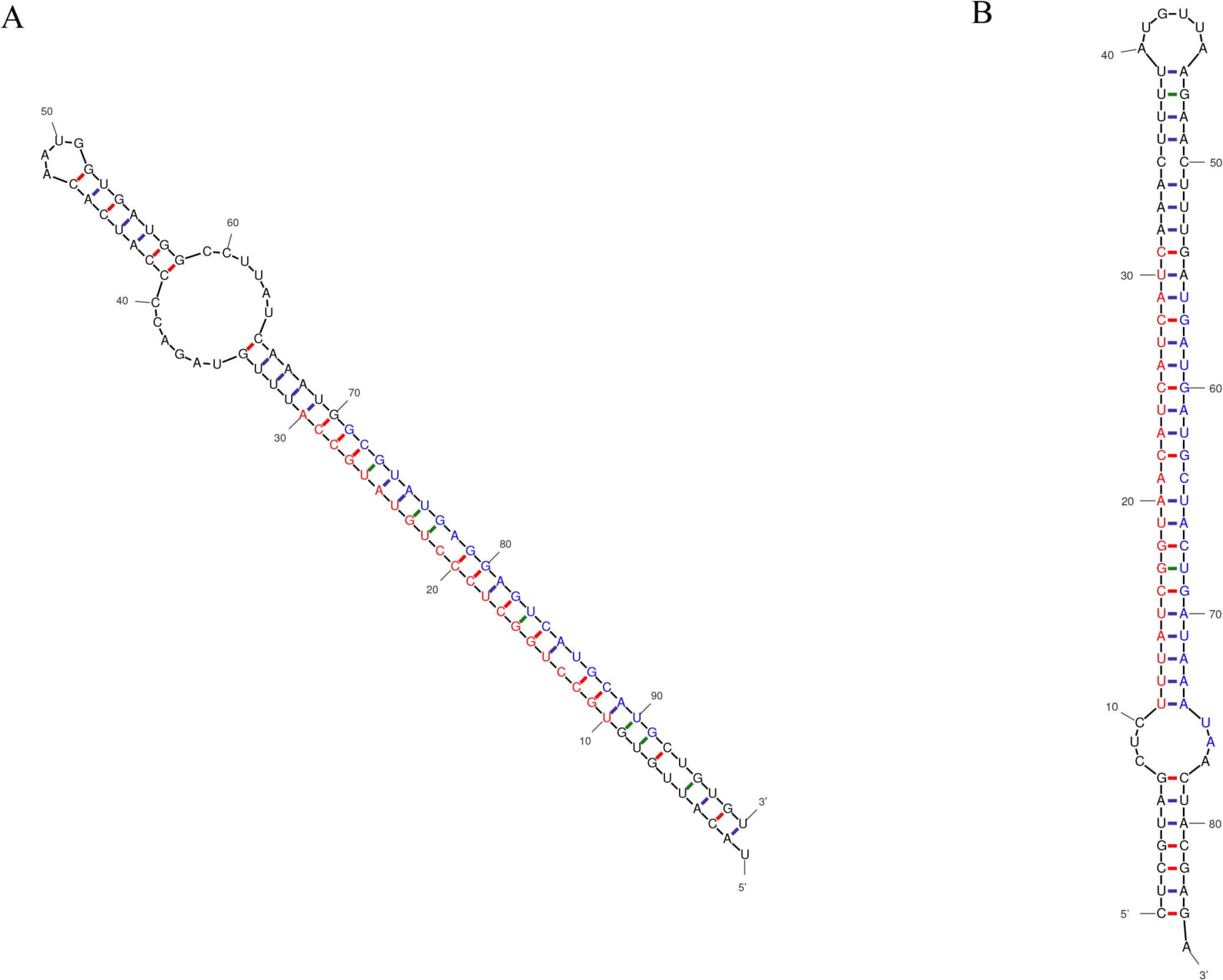
Predicted secondary structures of miRNAs. A, gma-MIR160f; B, novel_mir_003a. Red font represents miRNA-5p, and blue font represents miRNA-3p.

**Figure S2.**
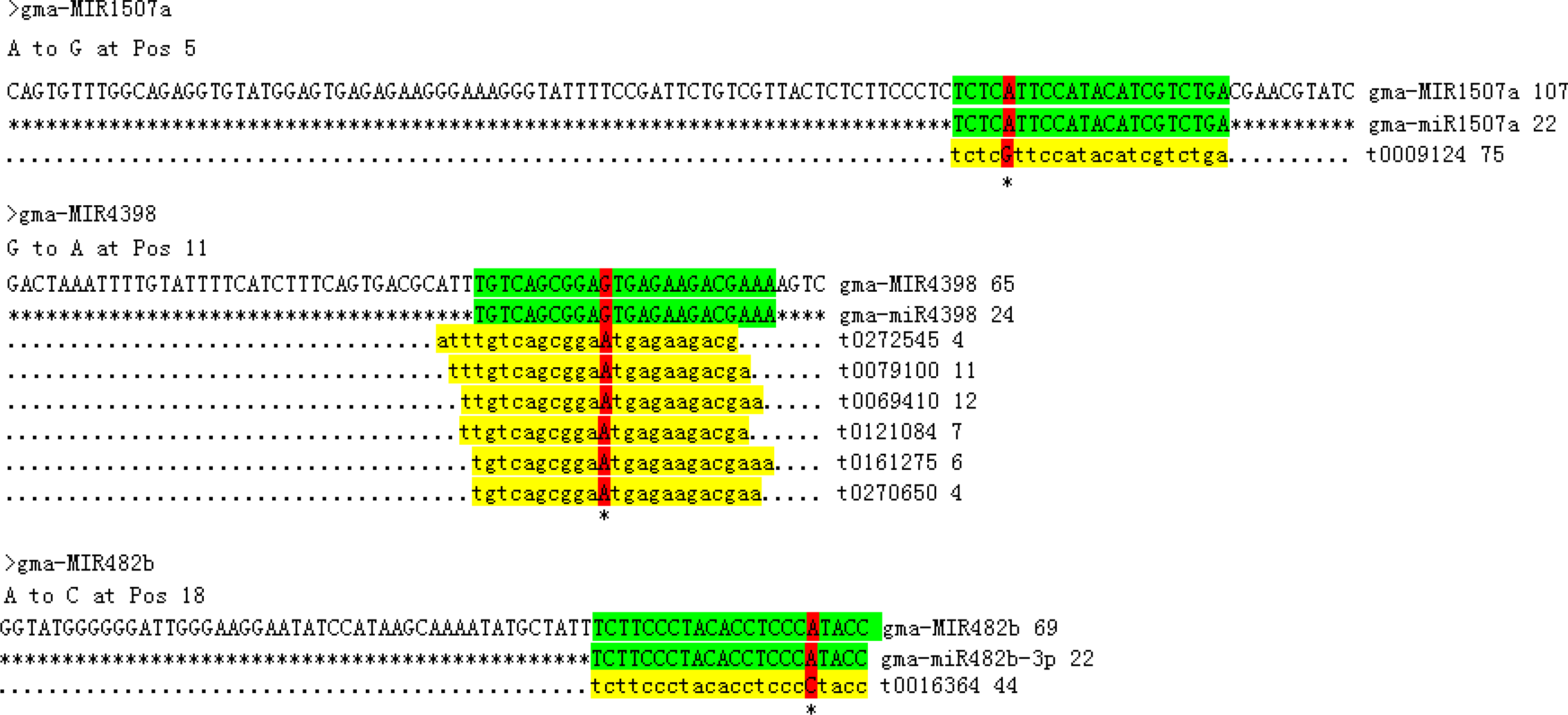
Expression profile of base-edited miRNA. The base-edited site is highlighted in red.

**Figure. S3.**
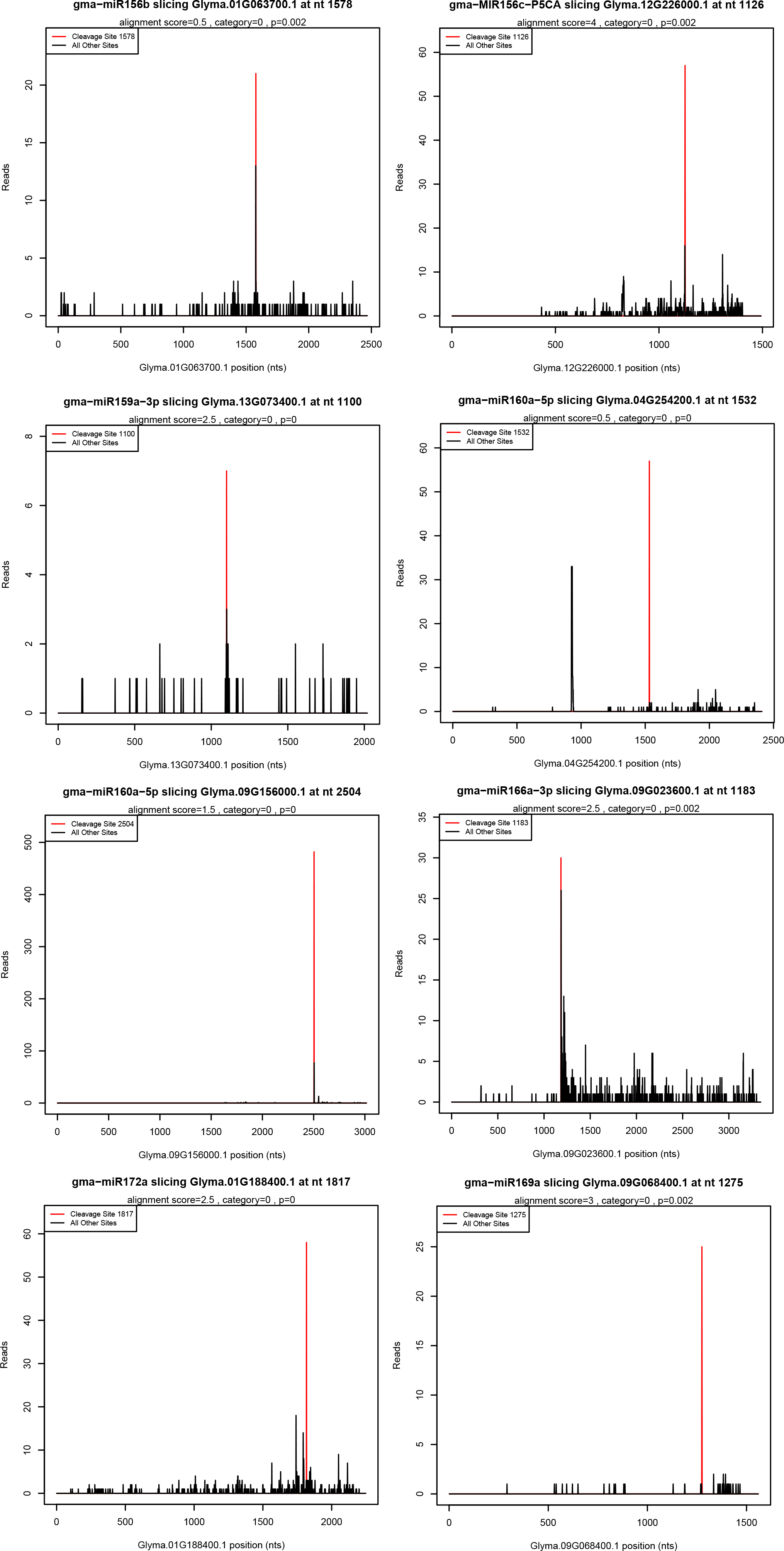
Target plots (t-plots) of identified miRNA targets. The X axis indicated the site position of target cDNA, the Y axis indicated the normal abundance of raw tags. The red line indicated the precited cleavage site.

**Figure S4.**
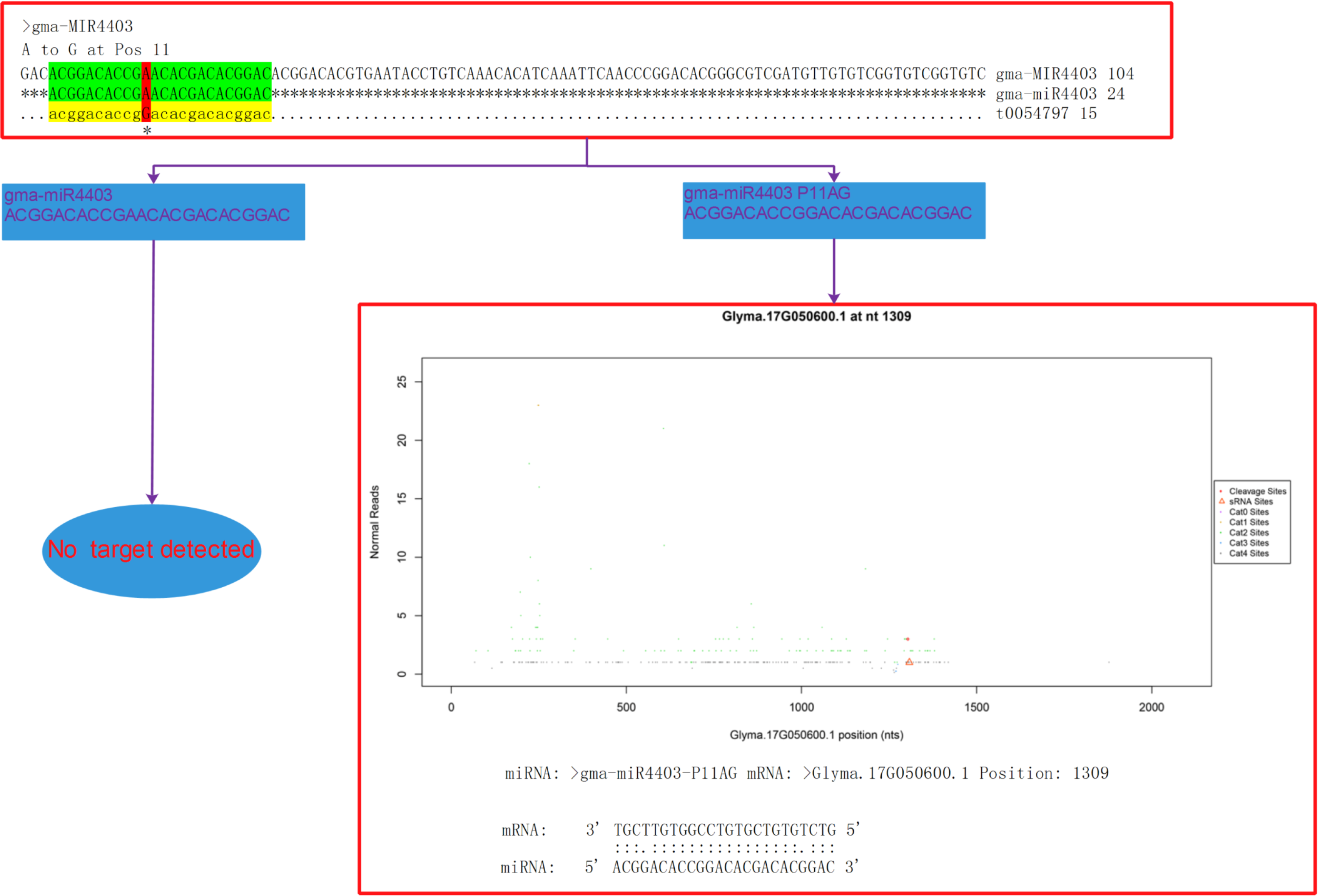
Targets identification of gma-miR4403 and its base edited miRNA.gma-miRA.

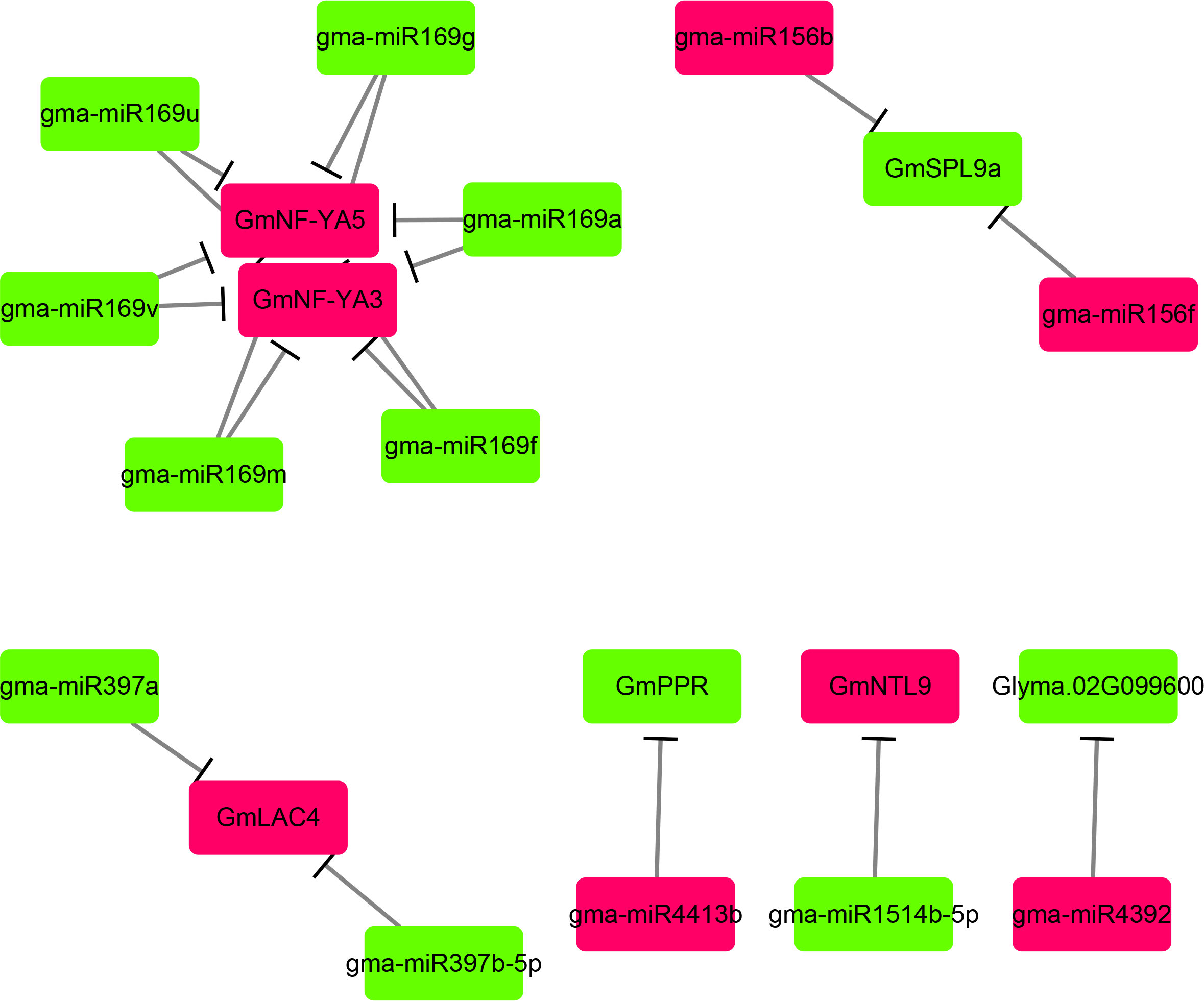

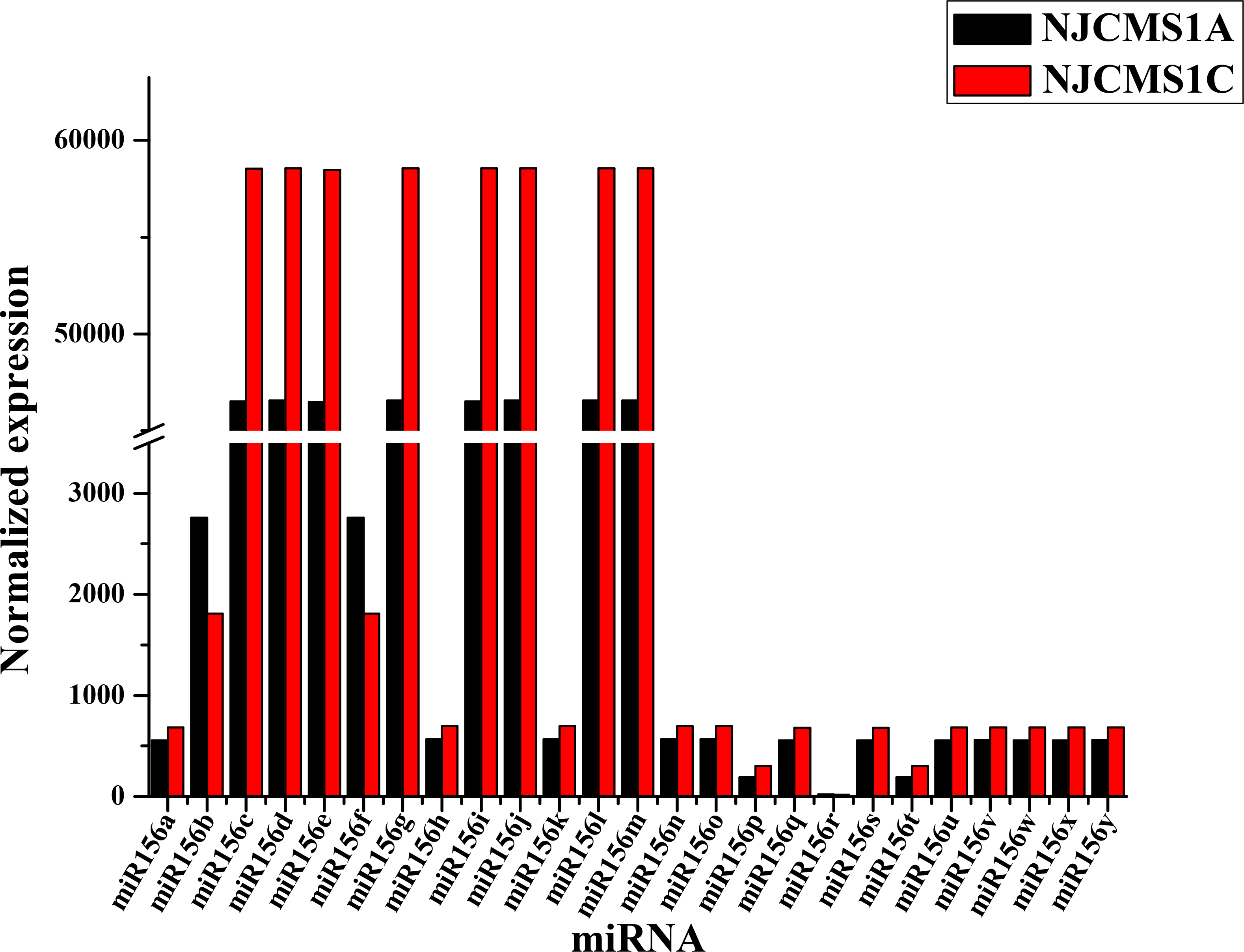

